# Investigating the genetic relationship of *Trichosporon asahii* isolates using a newly developed microsatellite typing assay

**DOI:** 10.1101/2024.11.12.623230

**Authors:** Elaine C. Francisco, Norma B. Fernández, Mauricio Carbia, Chendo Dieleman, Andra-Cristina Bostanaru-Iliescu, Jos Houbraken, Arnaldo L. Colombo, Ferry Hagen

## Abstract

The basidiomycetous yeast-like fungus *Trichosporon asahii* is increasingly encountered in the clinic and has been known to cause nosocomial infections. Here, we aim to develop a microsatellite typing tool to investigate the genetic relatedness of *Trichosporon asahii* isolates. Six microsatellite markers were selected from a nanopore long-read sequencing de novo assembled genome of the *T. asahii* type-strain CBS 2479. PCR reactions amplified three di-, tri-, or tetranucleotide repeats, respectively. All six microsatellite markers were used to analyze 111 *T. asahii* isolates, including single and related-patient isolates from clinical and environmental sources. Each marker exhibited between 11 and 37 alleles in this population, resulting in 71 different microsatellite genotypes. The Simpson’s diversity index ranged from 0.6452 to 0.8280 for the individual markers, and for the combined set of microsatellites 0.9793, indicating the high discriminatory power of the reported panel. In summary, this panel combines high reproducibility and specificity, making it suitable for use in epidemiological studies of *T. asahii* and in investigations of potential outbreaks caused by this pathogen.

## Introduction

*Trichosporon asahii* is an emerging yeast-like pathogen capable of causing life-threatening catheter-related infections worldwide (1–3). Despite being often overlooked, the occurrence of invasive trichosporonosis has dramatically increased in recent decades, with crude mortality rates reaching up to 80% depending on host comorbidities (3–6). Episodes of invasive trichosporonosis caused by *T. asahii* are primarily reported in long-term hospitalized patients with underlying hematological malignancies and neutropenia, as well as critically ill patients who have undergone invasive medical procedures, have indwelling medical devices, and have been exposure to broad-spectrum antimicrobial therapy (3,5). Recently, cases of invasive trichosporonosis have also been reported in immunocompetent hosts and hospitalized COVID-19 patients, posing new challenges in stratifying at-risk populations (7–11).

*Trichosporon asahii* exhibits a peculiar antifungal susceptibility profile, being intrinsically resistant to echinocandins and often showing decreased *in vitro* susceptible to amphotericin B (1,2,12,13), which can exert significant selective pressure on the growth of this pathogen. Triazoles, particularly voriconazole, are recommended as first-line therapy for treating invasive trichosporonosis (14). However, the intraspecific diversity among clinical *T. asahii* isolates may contribute to their reduced susceptibility to triazoles, highlighting the relevance of early diagnosis for effective management of invasive trichosporonosis (15,16).

While some authors have reported clusters of nosocomial *T. asahii* infections, there is a notable lack of epidemiological typing tools to investigate their potential clonal spread in clinical settings (11,17,18). Currently, sequencing of the intergenic spacer 1 (IGS1) of the ribosomal DNA (rDNA) is used to explore the intraspecific diversity of *T. asahii,* with 15 IGS1-genotypes already described (19). However, epidemiological studies assessing the global distribution of these genotypes have predominantly reported high prevalence rates for IGS-genotypes G1–G7. In contrast, the more recently identified IGS1-genotypes (G8–G15) have been documented mainly in single reports. This underscores the importance of employing a robust discriminatory typing approach to investigate the intraspecific diversity of clinical *T. asahii* isolates (15,16,20).

Microsatellites, also known as short tandem repeat units, are widely used as a fast, highly-sensitive, and cost-effective typing technique to investigate the molecular diversity within fungal populations during nosocomial outbreaks and for monitoring pathogens over time (21). Microsatellite-based typing has been recognized as the optimal tool for population studies and outbreak investigations in healthcare settings, particularly for *Candida auris*, *Candida parapsilosis*, *Aspergillus fumigatus* and *Cryptococcus* species (22–27), providing reliable evidence in the epidemiologic investigations. In this study, we developed a microsatellite-based typing for *T. asahii* and applied it to a large and genetically diverse collection of clinical and environmental isolates.

## Material and methods

### Media, strains, and standard DNA extraction

The clinical *Trichosporon asahii* type-strain CBS 2479 was used as a reference to set up the microsatellite typing panel, another set of 21 *T. asahii* isolates from the CBS culture collection (hosted by the Westerdijk Fungal Biodiversity Institute, Utrecht, The Netherlands) that originated from clinical (n=5), veterinary (n=5), and environmental (n=11) sources; a second large set of unique clinical isolates from Brazil (n=46), Argentina (n=9), Uruguay (n=4), and Romania (n=1). Additionally, a third set consisting of 29 sequential isolates, sampled on different days from 12 patients admitted to different medical centers in Brazil, and four isolates from Uruguay obtained from two patients was also used (Figure 1). Isolates were cultured onto malt extract agar and incubated for 48h at 25°C. DNA extraction was performed as described previously (28). All strains were identified as *T. asahii* based on sequencing the IGS1 rDNA locus, as previously described (15,20).

**Figure 1.**
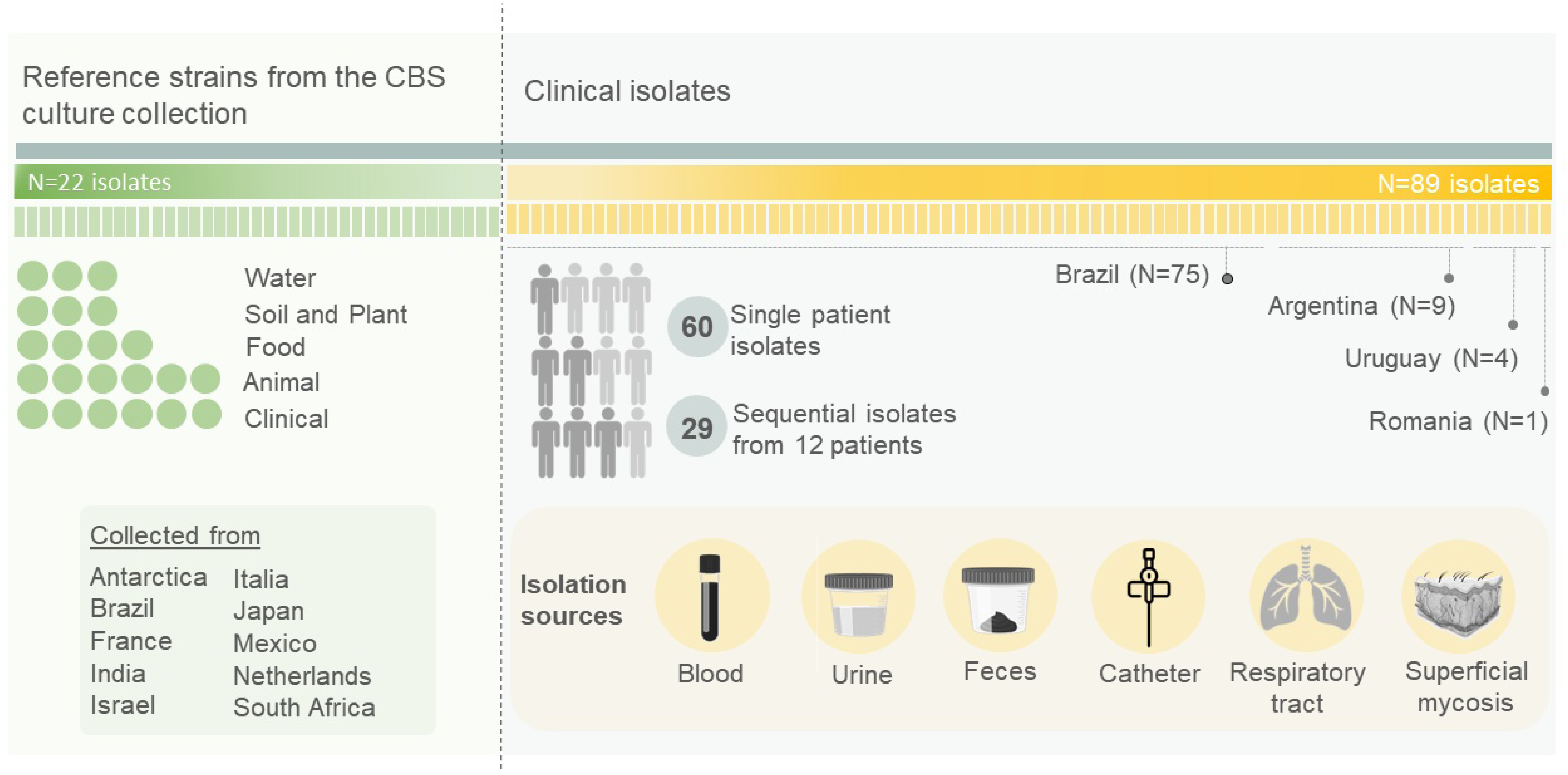
Graphical representation of the 111 *Trichosporon asahii* isolates evaluated in this study.

### Genome sequencing

The genome of the *T. asahii* type-strain CBS 2479 was published a decade ago and was at that time generated using short-read sequencing (NCBI BioSample SAMN02981437 and BioProject PRJNA164647). However, this draft genome was found to be highly fragmented with 78 scaffolds and 342 contigs (29). Hence, we re-sequenced the genome using long-read nanopore sequencing to cover complex genetic regions like the microsatellite loci. A fresh culture of CBS 2479 was prepared onto malt extract agar followed by 48h of incubation at 25°C, thereafter high-quality genomic DNA was extracted as previously in detail described by our team (30). Library preparation was performed using the native barcoding for genomic DNA kit (SQK-LSK109 in combination with EXP-NBD114; Oxford Nanopore Technologies, Oxford, UK) following the manufacturer’s instructions. The prepared library was first loaded onto a Flongle flow cell to confirm the quality, followed by running the library onto a MinION flow cell (both R9.4.1) and raw data were base called using Guppy v4.5.4 using the high-accuracy mode that yielded 4.8 Gbp of data with an N50 of 17 kbp and an N90 of 3.370 bp (all Oxford Nanopore Technologies). De novo genome assembly was performed using Flye v2.9. with the setting ‘--genome-size 24m--min-overlap 10000’ and resulted in a genome size of 25,250,028 bp dispersed over 19 fragments with an N50 of 2,305,946 bp and the largest fragment being 5,662,815 bp. The mean coverage was 198X for the nuclear fragment and 1,310X for the mitochondrial genome that had –after manual curation– a length of 31,421 bp. Data were deposited in NCBI with the accession numbers CP116781-CP116799 for the genome assembly, PRJNA926907 for the BioProject, SAMN32886118 for the BioSample, and SRR23205074 for the sequencing reads.

### Development of the microsatellite typing panel

The fasta-file of the de novo assembled CBS 2479 genome was used as input for the Tandem Repeat Finder software using the standard parameters, flanking regions for each locus were included to enable primer design (31). Close to 4,800 microsatellite loci were detected and were subjected to the following selection criteria: >10 copies of the repeat unit, >90% of the repeat units being intact, preferentially a di-, tri-, or tetranucleotide repeat unit, and selected loci had to be on different fragments of the de novo genome assembly of CBS 2479. This resulted in an initial list of 26 loci for which primers were developed using Primer3 v0.4.0 (32) with the standard settings that were only slightly adapted for: optimal primer T_m_ being 60°C ± 1°C, a maximum of 3 poly-X nucleotides and an optimal primer size of 20 bp (range 18–27 bp). The searched amplicon length was 50–200 bp, excluding the microsatellite loci.

First, a set of eight *T. asahii* isolates (CBS 2479, CBS 5599, CBS 7631, CBS 8969, L2122, L7918, L9206, and L920/2016), was used as the primary test set to test if the designed primer sets yielded amplicons for all isolates. PCRs were performed in a reaction containing 16.8 µl water, 2.5 µl 10× PCR buffer and 1.0 µl MgCl_2_ (50 mM), 1.0 µl 0.5 U BIOTAQ Taq polymerase (all Bioline, Meridian Bioscience, Memphis, TN, USA), 2.5 µl dNTP (1mM; Bioline), 0.1 µl 100 pmol/µl unlabeled forward and reversed primer (Integrated DNA Technologies, San Diego, CA, USA), and 1.0 µl DNA template. PCR was performed as follows: initial denaturation at 94 °C for 5min, 35 cycles of 94 °C for 30 sec, 60 °C for 30 sec, and 72 °C for 1 min, a final extension for 72 °C for 5 min and hold at 21 °C. To check the success of the designed primer sets all amplicons were checked by 2% agarose gel electrophoresis.

Twelve out of the 26 initial primer sets yielded an amplicon for the set of 8 test isolates, these twelve primer sets were subsequently tested using a 2^nd^ larger set of an additional 16 *T. asahii* isolates that went through the same PCR procedure as described above. Six primer combinations remained that yielded amplicons for all 24 isolates tested as well as size differences were observed by agarose gel electrophoresis, for each of the primer sets a primer with a fluorescein-label was ordered to enable the detection of amplicons by capillary electrophoresis (Table 1).

**Table 1.**
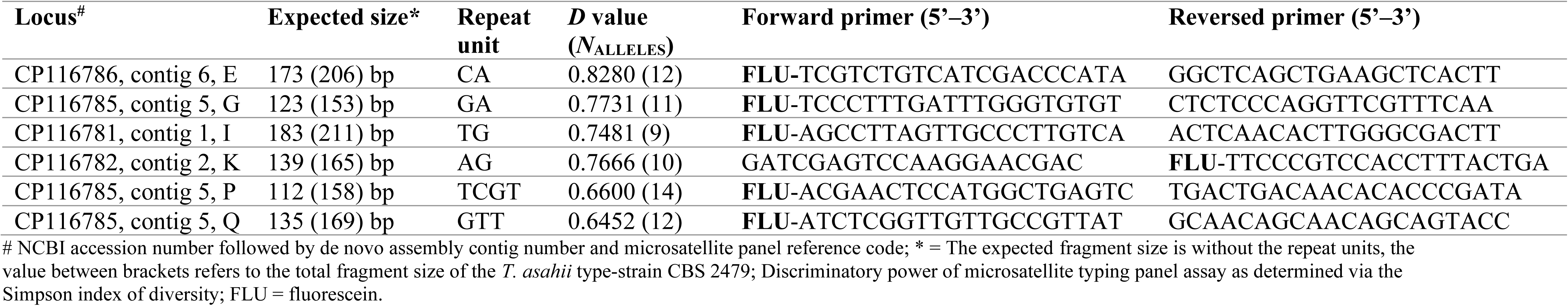
Microsatellite typing panel details.

Finally, a large set of 75 clinical isolates obtained from South America and Europe was used to check for reproducibility, stability, and specificity of the *T. asahii* microsatellite typing panel of six loci developed. The isolates were obtained from different anatomical sites, including deep-seated and superficial infections, representing the five most prevalent IGS1-genotypes as previously described (15).

For capillary-based fragment analyzes the PCR approach as described above was followed, however, the six PCRs were performed with a fluorescein-labeled primer (Table 1). PCRs were checked by 2% agarose gel electrophoresis for their yields. Thereafter, amplicons were Sephadex-purified (Sigma-Aldrich, Darmstadt, Germany), and amplicons were arbitrarily 50–200× diluted with water. One microliter diluted amplicon was mixed with 10× diluted Orange500 size marker (Nimagen, Nijmegen, The Netherlands) in a 96-well plate followed by 1 min 94 °C and 1 min 4 °C. Raw data was obtained by running the fragment analysis onto an ABI3700xL Genetic Analyzer (Applied Biosystems, Palo Alto, CA, USA).

### Data analysis and discriminatory power

Raw data and relatedness between strains were analyzed using Bionumerics v7.6 (Applied Maths, Sint-Martens-Latem, Belgium) via the unweighted pair group method with arithmetic averages (UPGMA) as previously described (24). The discriminatory power of the microsatellite panel was determined using the Simpson index of diversity (*D*) (33). A *D*-value of 1 indicates that the typing method was able to discriminate between all isolates, while a value of 0 indicates that all isolates were identical (clonal).

## Results

### Development of T. asahii microsatellite typing assay

The microsatellite typing panel was developed based on a selection of six out of 26 promising loci, being four di-nucleotide (loci E, G, I and K), one tri-nucleotide (locus Q) and one tetra-nucleotide repeat loci (locus P). Locus G, P and Q were all located on the same contig, while E, I and K were on different contigs (Table 1).

### Evaluation of T. asahii microsatellite typing panel in a diverse collection of clinical isolates

A set of 111 *T. asahii* isolates, including 22 CBS reference strains, 60 non-replicated single patient, and 29 replicated clinical isolates from 12 patients, was used. The Simpson index of diversity (*D*) ranged from 0.6452 (locus Q, tetra-nucleotide repeat unit) to 0.8280 (locus E, di-nucleotide repeat unit) (Table 1). The combination of all six loci yielded a *D* value of 0.9793.

Among the 111 *T. asahii* isolates tested, we identified 71 microsatellite genotypes, which each genotype clustering 1 to 11 isolates (Figure 2). Of the 46 non-replicated Brazilian clinical isolates, 20 exhibited unique STR markers (44%). Notably, four Uruguayan isolates collected from two different patients (Patient 1 = MC215 and MC216; Patient 2 = MC217 and MC218) were identical, and one Brazilian isolate exhibited the same genotype. Among the nine Argentinian *T. asahii* isolates evaluated, six displayed unique STR markers, two shared identical genotypes, and one clustered in a separate group being hitchhiking with isolates collected from both clinical and environmental sources.

**Figure 2.**
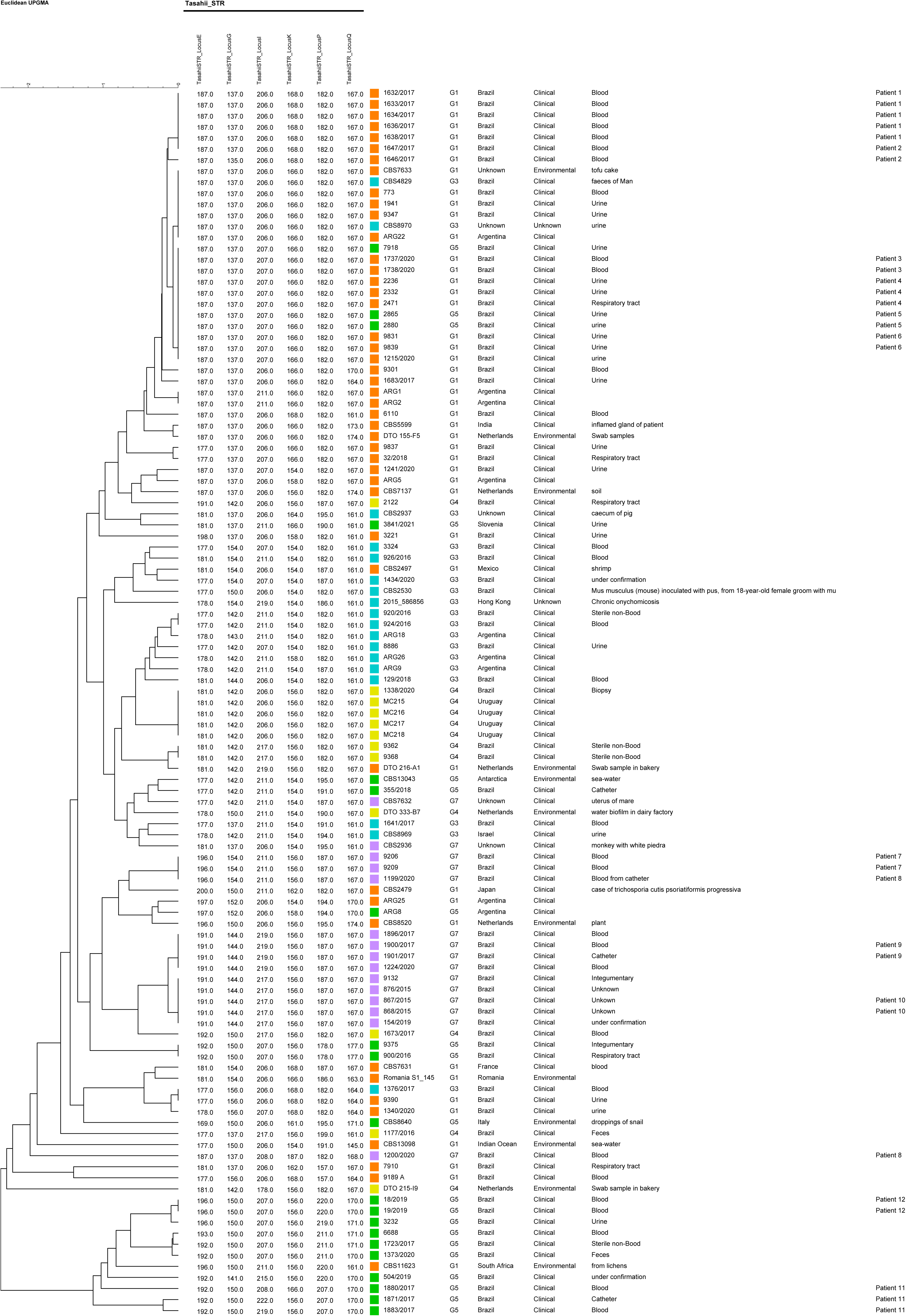
UPGMA dendrogram showing genotypic diversity among 111 *Trichosporon asahii* isolates obtained from clinical and environmental samples. The scale bar indicates the percentage similarity between the genotypes. The dendrogram was generated based on analysis of six microsatelite markers.

The 29 sequential clinical isolates collected from 12 patients were categorized into 11 genotypes. Sequential isolates from three patients showed distinct genotypes, grouping into different microsatellite clusters, as follows: isolates from patient 2 (L1646/2017 and L1647/2017) exhibited a single difference at locus G; isolates from patient 8 (L1199/2020 and L1200/2020) displayed varying numbers of microsatellite repeat units across all six loci examined; while isolates from patient 11 (L1871/2017, 1880/2017, and 1881/2017) differed at loci I and K; The sequential isolates from the remaining patients clustered with their sequential counterparts based on microsatellite genotypes (Figure 2).

### IGS1 sequence-based genotyping versus microsatellite typing

The neighbor-joining method divided the 111 *T. asahii* isolates into 5 IGS1-genotypes (G1 n=47, 42.3%; G3 n=17, 15.3%; G4 n=12, 10.8%; G5 n=20, 18%; and G7 n=15, 13.5%). Brazilian clinical isolates were represented by all these five IGS1-genotypes. Majority of the Brazilian isolates were IGS1-genotype G1 (n=30, 40%), followed by G5 (n=16, 21.3%), G7 (n=13, 17.3%), G3 (n=10, 13.3%), and G4 (n=6, 8%).

Even though the isolates sharing identical IGS1-genotypes, such as the four clinical isolates from Uruguay (G4), non-concordance results between IGS1 sequencing-based genotypes and microsatellite typing were observed (Figure 3A and 3B; Supplementary Figure 1).

**Figure 3.**
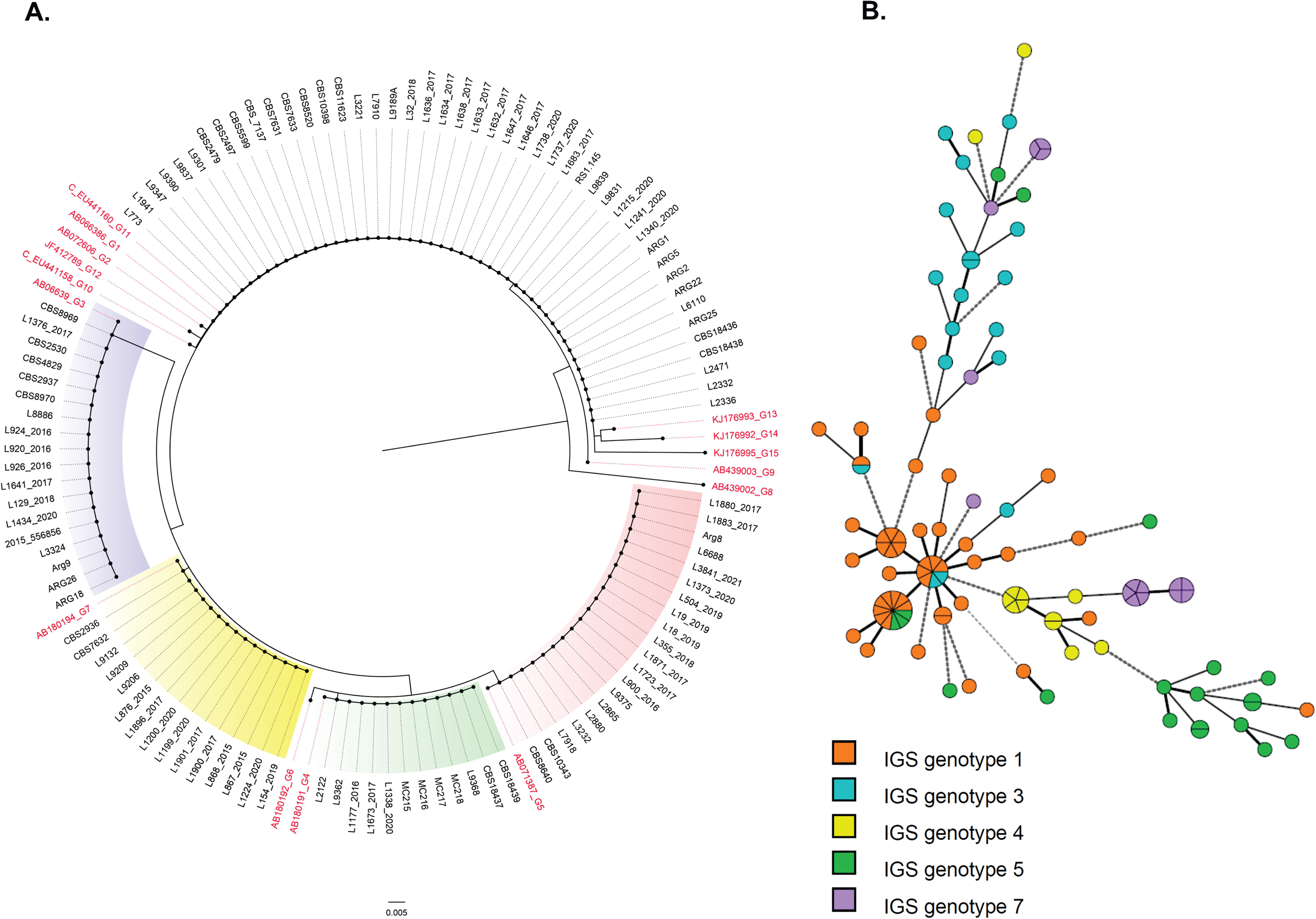
Comparisons between IGS1-sequence-based genotyping and the developed microsatellite panel of 111 *T. asahii* isolates. A: Neighbor-joining tree based on the rDNA sequencing target. Reference strains of the described genotypes are shown highlighted in red corresponding to their NCBI GenBank accession numbers. B: minimum spanning tree (bottom panel) generated by six loci of microsatellite data showing the relationship between microsatellite typing and IGS1-genotypes. Each circle represents a unique genotype, if multiple isolates share an identical genotype, they are shown as a fraction of the circle. Lines between circles indicate the relative similarity, the shorter the line the less prominent was the difference between the microsatellite genotypes. Thick lines identify genotypes with fewer differences (one locus out of six tested), while medium thick lines identify genotypes that have two out of the six loci different, thick and medium thick dashed lines represent genotypes that share 3 or 2 identical loci out of six tested. Thin dotted lines identify genotypes sharing only one identical locus.

### Reproducibility and specificity of the developed microsatellite typing panel

To test the reproducibility of the developed panel, the first set of eight isolates (described above) were independently amplified at four least times. In all replicate assays, the microsatellite markers showed identical profiles for all isolates evaluated, indicating reproducible results. Finally, the validation of the developed typing panel for *T. asahii* was performed by testing 20 *Trichosporon* isolates (representing the 11 different non-*T. asahii* species currently recognized to the genus), added to representative isolates of the correlated genera *Apiotrichum* and *Cutaneotrichosporon*. PCR product amplification for all the six selected loci was not found demonstrating its highly specie-specific typing toll for *T. asahii*.

## Discussion

Here, we report the development of the first microsatellite typing panel to genotype *T. asahii*. This basidiomycete yeast-like pathogen is able to cause a broad spectrum of human infections that have attracted medical attention due to its increasing prevalence in life-threatening infections worldwide (3,5,12,14). Even recognized as the second/third non-*Candida-*related yeast obtained from invasive infection episodes, it competes with *Cryptococcus* as leading basidiomycetous yeast pathogen. At present, IGS1 sequence-based genotyping is the preferred approach to investigate genetic diversity among *T. asahii* isolates (3,15,34). Despite its relevance, the IGS1-based genotyping lacks the genetic diversity to use it as an outbreak and epidemiological typing tool which makes it difficult to use it for nosocomial outbreak investigation.

Microsatellites, short repetitive DNA sequences, are widely employed in molecular studies to study the genetic relatedness between isolates within fungal populations. This typing tool has significantly contributed in the advancement of epidemiological typing for a variety of human pathogens, including *Candida auris*, *Candida parapsilosis*, *Nakaseomyces glabratus* (syn. *C. glabrata*), *Cryptococcus neoformans*, *Cryptococcus deneoformans*, *Cryptococcus deuterogattii* and *Aspergillus* species (25–27,34–39). Compared to other DNA-typing tools, like ITS-based and amplified fragment length polymorphism (AFLP) profiling, microsatellites have been demonstrated to have superior performance in epidemiological studies (23,24,40,41).

The here described *T. asahii* microsatellite typing panel, that consists of six loci, revealed a remarkable genetic diversity among *T. asahii* isolates. This novel panel successfully distinguished 82 unrelated isolates (comprising 22 CBS reference isolates and 60 clinical isolates) into 58 distinct genotypes, a significantly better genetic discriminatory power compared to the gold standard of IGS1-genotyping that identified five IGS1-genotypes in the same isolate set. Notably, our panel effectively differentiated between isolates based on their origins, anatomical sites, as well as antifungal fungal susceptibility, and year of isolation. Moreover, 100% similarity was observed among sequential isolates obtained from 9 of the 12 patients from which multiple isolates were available. Sequential isolates from the remaining three patients were distributed across different microsatellite genotypes, all of these sequential isolates were collected two to seven days after the first isolate, suggesting potential co-infection of these patients by different *T. asahii* genotypes. To underscore he limitation of IGS1-genotyping, all these sequential isolates shared the same IGS1-genotype.

Interestingly, while most isolates clustered with their sequential counterparts based on microsatellite profiles, suggesting stable genetic lineages, the observed variations indicate underlying biological complexities. These discrepancies could arise from factors such as phenotypic plasticity, environmental adaptation, or genetic mutations not captured by microsatellite analysis. Further investigation into additional biological and environmental characteristics could provide insights into the adaptive mechanisms and epidemiological dynamics of *T. asahii*.

At present, there is a lack of studies comparing different typing tools for *T. asahii*. IGS1-sequencing is a powerful tool and is even nowadays considered the gold standard in reference laboratories globally to discriminate the *Trichosporonales* genera and species, and even up to the different genetic lineages within *T. asahii* (20). This was recently nicely presented by Francisco and co-workers (2019) who performed molecular characterization of a set of locally collected *Trichosporon* species isolates (2). This endeavor led to the identification by them of a genetic lineage that we know since recently as *T. austroamericanum* (42). However, from the various epidemiological studies, it can be concluded that IGS1-sequencing is lacking the needed resolution for outbreak typing (15,16,20,43–45).

To understand the genetic relationship between *T. asahii* isolates from elderly patients hospitalized in a single Spanish care center, Treviño and team (2014) supplemented IGS1-sequencing with the (nowadays discontinued) commercial DiversiLab typing tool (46). However, the addition of that rep-PCR typing tool led to inconclusive results, as the fingerprint patterns lacked sufficient discriminatory power. The genome-wide based typing tool Amplified Fragment Length Polymorphisms (AFLP) analysis has been shown to be an informative approach for fungal outbreak investigations, it has been reported only once for *Trichosporon* (47). Unfortunately, the selective primer combination used by Ahangarkani and colleagues (2021) resulted in AFLP profiles that could not distinguish potential related isolates to the unrelated ones. Moreover, AFLP genotyping has been found to be more laborious, costly, and less reproducible than other typing tools, such as microsatellite typing. Hence, the latter has gradually replaced the former to investigate outbreaks caused by fungal pathogens.

Recently, Pumeesat & Wongsuk (2023) published a multi-locus sequence typing (MLST) assay consisting of sequencing five nuclear loci that they applied onto a set of 51 clinical Thai *T. asahii* isolates. Although these authors concluded that their MLST was useful for population structure analysis it seems to have limited genetic diversity to use it for outbreak investigations as the 51 isolates were dispersed among only 5 sequence types (48). Desnos-Ollivier and colleagues (2020) used whole genome sequencing on a subset of 32 out of 54 *T. asahii* isolates that were collected during a ∼17-years period. Initially, these isolates were typed using IGS1-sequencing and it is therefore not surprising that short-read genome sequencing resulted in a higher discriminatory power compared to the former typing approach (49). In the foreseeable future genome sequencing will get a more prominent role in investigating fungal nosocomial outbreaks. However, this approach is currently and, in the years ahead, a too costly typing tool for diagnostic laboratories in low-and middle-income countries. Thus, an intermediate high-resolution typing approach for these laboratories will be the microsatellite typing scheme that we here presented to rapidly investigate genetic relatedness of *T. asahii* potentially involved in nosocomial outbreaks.

In conclusion, we developed a fast, reproducible, and species-specific microsatellite typing assay for *T. asahii* able to screen many isolates for epidemiological typing purposes, transmission routes, and outbreak investigations.

## Statements Ethical approval

This study was approved by the Research Ethics Committee of Universidade Federal de São Paulo (CEP-UNIFESP 6183240519/2019).

## Supplementary files

**Supplementary Figure 1.** The phylogenetic analyses of 111 *T. asahii* isolates and the IGS1 genotypes was inferred using the Neighbor-Joining method. The optimal tree is shown. The percentage of replicate trees in which the associated taxa clustered together in the bootstrap test (1000 replicates) are shown next to the branches. The tree is drawn to scale, with branch lengths in the same units as those of the evolutionary distances used to infer the phylogenetic tree. The evolutionary distances were computed using the Kimura 2-parameter method and are in the units of the number of base substitutions per site. This analysis involved 126 nucleotide sequences, conducted in MEGA X.

