## Supplementary figures and images for "Investigating the genetic relationship of *Trichosporon asahii* isolates using a newly developed microsatellite typing assay"

### Supplemental Figure 1

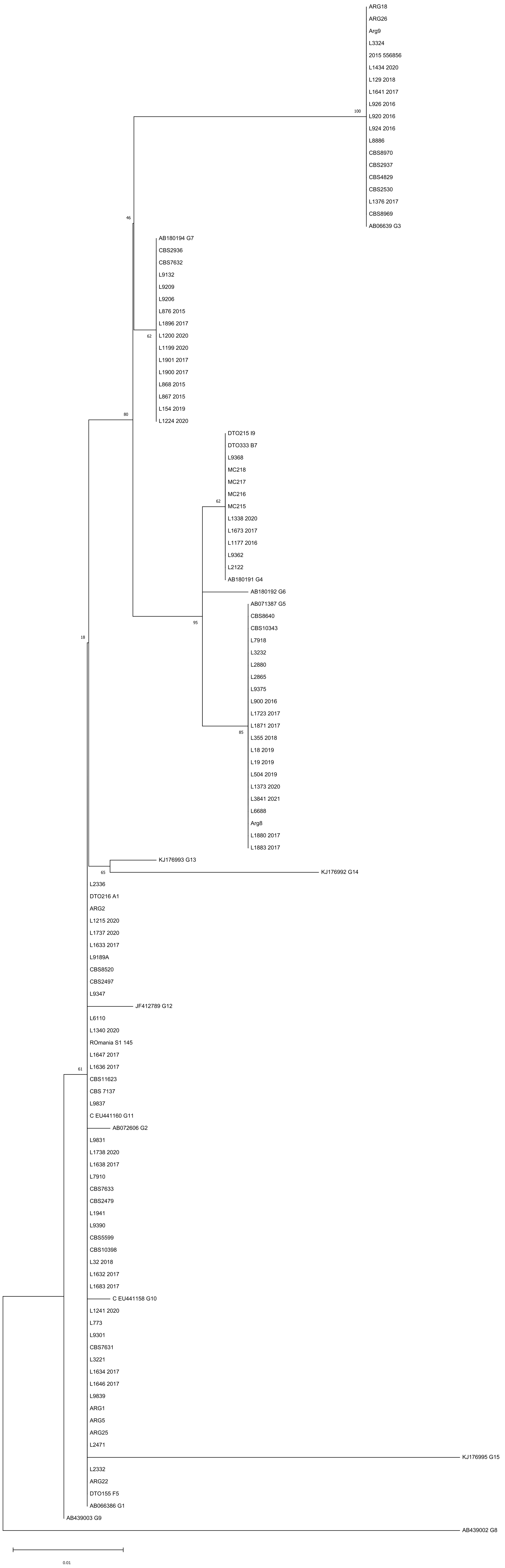
